# Hungry for compliments? Ghrelin is not associated with neural responses to social rewards or their pleasantness

**DOI:** 10.1101/2022.11.18.516889

**Authors:** Uta Sailer, Federica Riva, Jana Lieberz, Daniel Campbell-Meiklejohn, Dirk Scheele, Daniela M. Pfabigan

## Abstract

The stomach-derived hormone ghrelin motivates food search and stimulates food consumption, with highest plasma concentrations before a meal and lowest shortly after. However, ghrelin also appears to affect the value of non-food rewards such as interaction with rat conspecifics, and monetary rewards in humans. The present pre-registered study investigated how nutritional state and ghrelin concentrations are related to the subjective and neural responses to social and non-social rewards.

In a cross-over feed-and-fast design, 67 healthy volunteers (20 women) underwent functional magnetic resonance imaging (fMRI) in a hungry state and after a meal with repeated plasma ghrelin measurements. In task 1, participants received social rewards in the form of approving expert feedback, or non-social computer reward. In task 2, participants rated the pleasantness of compliments and neutral statements.

Nutritional state and ghrelin concentrations did not affect the response to social reward in task 1. In contrast, ventromedial prefrontal cortical activation to non-social rewards was reduced when the meal strongly suppressed ghrelin. In task 2, fasting increased activation in the right ventral striatum during all statements, but ghrelin concentrations were neither associated with brain activation nor with experienced pleasantness. Complementary Bayesian analyses provided moderate evidence for a lack of correlation between ghrelin concentrations and behavioral and neural responses to social rewards, but moderate evidence for an association between ghrelin and non-social rewards.

This suggests that ghrelin’s influence may be restricted to non-social rewards. Social rewards implemented via social recognition and affirmation may be too abstract and complex to be susceptible to ghrelin’s influence. In contrast, the non-social reward was associated with the expectation of a material object that was handed out after the experiment. This may indicate that ghrelin might be involved in anticipatory rather than consummatory phases of reward.

## 1. Introduction

Ghrelin is an appetite-stimulating hormone produced by the stomach (1). Ghrelin concentrations increase during energy restriction (2,3) and decrease following eating (4,5). In humans, ghrelin concentrations correlate with feelings of subjective hunger (e.g.; 6,7). Ghrelin regulates food-related behaviours (for a review, see 8), but it is now also established that its functions go far beyond its role as an appetite stimulant.

By acting on reward-processing areas in the brain, ghrelin can influence both general and specific motivation to receive rewards. For instance, administration of ghrelin increased alcohol intake in mice (9) and the motivation to self-administer heroin in rats (10). In humans, ghrelin concentrations correlated with craving in alcohol-dependent individuals (11,12), with the pleasantness of odours (13) and reward system activation to food pictures (14). Furthermore, the suppression of ghrelin signalling reduced sexual motivation (15) and the rewarding effects of amphetamine and nicotine in mice (16,17). These effects are thought to occur via ghrelin’s effect on dopaminergic functions in the ventral tegmental area (VTA) (18,19) from where dopaminergic neurons project to the ventral striatum. As such, ghrelin may modulate neural reward responses to stimuli other than food.

Indeed, ghrelin seems to modulate the response to monetary rewards in humans. Lower ghrelin concentrations were associated with slower choices following monetary rewards and increased reward-related activity in dorsolateral prefrontal cortex in obese participants (20). In a different study, healthy individuals and individuals with a low-weight eating disorder were asked to choose between smaller immediate and larger delayed monetary rewards. Only in healthy participants, higher ghrelin concentrations predicted a stronger preference for smaller immediate over greater delayed monetary rewards (21). Moreover, gambling cues in a laboratory casino setting increased ghrelin levels particularly in fasted participants, and these ghrelin increases predicted gambling persistence when participants were confronted with continued monetary losses (22). Ghrelin concentrations have also been associated with a general motivation to approach goal-oriented outcomes as measured with a questionnaire (23).

Recent research indicates that ghrelin might also play a role in social reward. Ghrelin modulated the preference for social interaction in rats (24). In this study, ghrelin increased and a ghrelin antagonist decreased the preference for social interaction, although only in the larger rat in a pair. Ghrelin receptor signalling is also associated with social motivation in mice: Mice lacking the ghrelin receptor or treated with a ghrelin antagonist approached conspecifics with longer latencies and spent less time in interaction (25). First evidence in humans also hints at a potential social role of ghrelin: Following a meal, lonelier women showed a ghrelin response that differed from that of less lonely women (26). Moreover, lower reward-related activation in the orbitofrontal cortex was observed in response to social interaction in the form of touch when ghrelin concentrations were high (27).

Based on these previous findings, we investigated the role of ghrelin in the neural processing of social and non-social rewards in humans in the current study. Specifically, we used functional magnetic resonance imaging (fMRI) to scan healthy volunteers while they received indirect social as well as non-social reward (task 1), and direct social reward (task 2). The social reward consisted of an endorsement of music preferences by experts in task 1 and of social affirmation in the form of compliments in task 2. Participants rated also the pleasantness of these compliments. Non-social reward was provided in task 1 in the form of computer algorithm feedback, which then resulted in an USB stick with participants’ favourite songs for them to take home. Participants were invited twice to the laboratory in a fasted state equivalent to high ghrelin concentrations. In one session, participants stayed fasted (no-meal session) to maintain high ghrelin concentrations, while in the other session they received a standardized meal to decrease ghrelin concentrations (liquid-meal session).

Given evidence that ghrelin affects the response to various types of social and non-social rewards in rodents and humans, we hypothesized that it may also modulate the processing of these rewards in humans. We expected ghrelin concentrations (i.e. concentration differences following a meal versus no meal) to correlate with activation differences between meal versus no meal in key areas of the brain’s reward circuit (e.g., bilateral ventral striatum, ventromedial prefrontal cortex, bilateral orbitofrontal cortex), as well as with meal versus no meal differences of behavioural outcomes in both tasks. Our preregistered bi-directional hypothesis stated that ghrelin could either have reward-enhancing (9,10,15,16,24) or reward-diminishing effects (28).

## 2. Methods

The study was preregistered on the Open Science Forum (https://osf.io/f9rkq). Whenever the procedure and/or the analyses are deviating from the pre-registration, it is clearly stated.

### 2.1 Participants

The pre-registered number of participants was 80, based on a power analysis performed with PANGEA (29) for means-based within-subject analyses of variance (ANOVAs): power > 80%, a medium effect size of d=0.45 for the highest interaction effect, one replication per cell, calculated for behavioural effects in each imaging task. We also took a drop-out rate of about 15% into account. For the social-recognition-by-experts task (task 1) with a 2 x 2 x 2 within-subjects design (nutritional state x reward type x outcome), at least 40 participants were recommended to reach the intended power. For the social affirmation task (task 2) with a 2 x 2 within-subjects design (nutritional state x statement type), at least 67 participants were recommended.

The COVID-19 pandemic did not allow us to reach the planned sample size though. However, we changed the analysis plan and conducted a single-trial analysis with linear mixed models instead of a means-based ANOVA design for the behavioural data of task 2. This new analysis reaches a power of 80% already with 16 participants (calculated post-hoc with PANGEA).

Initially, 68 participants between 18-55 years of age (47 men, mean age: 27.8, SD: 7.8; 20 women, mean age: 31.9, SD: 10; and one participant who did not wish to disclose their sex/gender, age: 30) were recruited via flyers and social media. Inclusion criteria were Norwegian language skills, a body mass index (BMI) between 18 and 29.9 kg/m^2^, normal or corrected to normal vision, and fulfilment of MRI safety criteria. In addition, participants were required to report no history of eating disorders, diabetes, gastrointestinal surgery, to tolerate lactose, to not suffer from any mental disorders at the time of the experiment, and to not take any drugs affecting gastrointestinal function. Female participants were required not to take hormonal contraceptives and to conduct the experiment in the first week of their menstrual cycle (between days 1-8) in order to reduce potential influences of sex hormones on plasma ghrelin concentrations (30) or on reward processes (31–33).

One male participant dropped out after the first blood sample because of circulation problems. Seven participants were not available for the second session that was delayed due to a Covid-19 lockdown. Thus, 60 participants (42 men, 17 women, 1 undisclosed individual) attended both measurement session. Due to technical problems during scanning, the number of available participants per analysis is different in tasks 1 and 2, and will be explicitly stated in the results section.

Handedness was assessed using the Edinburgh Handedness Inventory (34). The majority of participants was right-handed (n=53) and of self-declared European descent (n=66). Five participants were left-handed, eight were ambidextrous (n=2 missing); and two of self-declared Asian descent.

Participation was reimbursed with universal gift cards of ~ EUR 150 per person for both sessions lasting four to five hours each. The session contained further experimental tasks which were (27,35) and will be presented elsewhere. Written informed consent was obtained from all participants prior to the experiment. The study was conducted in accordance with the latest version of the Declaration of Helsinki and approved by the Regional Committee for Medical and Health Research Ethics (REK South-East B, project 26699).

### 2.2 Procedure

The experiment took place at the Institute of Basic Medical Sciences, University of Oslo. Each participant was scheduled for two sessions on separate days – see Figure 1 for a graphical overview of the experimental procedure. In order to change endogenous ghrelin concentrations, participants received a standard liquid meal (described further down) at the beginning of one session (liquid-meal session), but not the other (no-meal session). The aim was to decrease ghrelin concentrations with the liquid meal, and to keep them at a high level without the meal (36). Session order was pseudo-randomized across participants, and no-meal/liquid-meal sessions were by median four days apart (range 1-85). In total, data from participants in 65 liquid-meal and 62 no-meal sessions were collected.

**Figure 1:**
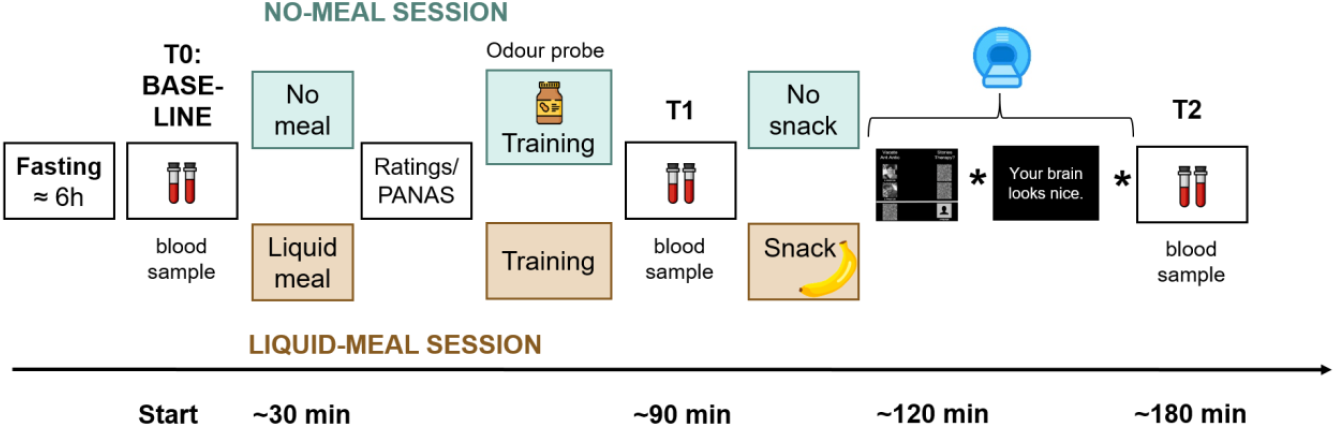
Sequence of experimental procedures. Participants arrived fasted. A baseline collection of a blood sample and a blood glucose measurement were performed (time point T0/Baseline). In one session, a liquid meal was provided before the tasks (apricot colour), in the other participants remained fasted (mint colour). After consumption of the liquid meal and at a corresponding time point in the no-meal session, participants rated their current bodily and affective states. Afterwards, participants performed several training trials on a laptop to familiarise them with the tasks. Only in the no-meal session, participants were presented with an odour probe. Then, another blood sample was collected (T1). Shortly before scanning, a snack was provided in the liquid-meal session only. The stars in the time line represent placeholders for two further experimental tasks that were conducted in the scanner and will be presented elsewhere. After scanning, a third blood sample was collected (T2).

The experiments were either scheduled at 3 pm or 4.30 pm. Participants were required to fast at least six hours prior to the experiment and drink no more than one liter of water during that time. To increase participants’ compliance with fasting, they were informed that blood glucose concentrations would be checked at the beginning of each test. For this, a pin prick test was conducted with an Accu-Check^®^ Aviva device (Roche Diagnostics Norge AS).

Afterwards, the first blood sample was taken (T0/Baseline). Saliva samples were also taken, the results of which are reported elsewhere (27). Body composition parameters such as weight, fat mass, and adipose visceral tissue were measured with a bioelectrical impedance device (seca mBCA 515, seca gmbh & co. kg., Germany) always during the second test session. The standardised meal consisted of 300 ml of raspberry flavoured, fat-free, fermented milk (Biola^®^, Tine BA; 50 kcal/100 g) and 300 ml of chocolate milk (Sjokomelk, Tine BA; 58 kcal/100 g). Participants consumed this meal within 15 minutes after the first blood sample in the liquid-meal session. In contrast, in the no-meal session, they stayed fasted and were offered the two beverages only at the end of the experiment.

After the meal in the liquid-meal session and at a corresponding time point in the no-meal session, participants rated their current bodily states via an online tool of the University of Oslo called nettskjema. Using a five-point Likert scale (1=not at all to 5=very much), answers were collected on subjective hunger feelings (“How hungry are you right now?”), feelings of stomach emptiness (“How full is your stomach right now?”), subjective thirst (“How thirsty are you right now?”), desire to eat (“How much do you want to eat food right now?”), and the estimated amount participants could eat (“How much food could you eat right now?”). They also answered one question on how much they would be willing to pay (in NOK) for their favourite food right now, and how much time had passed since their last meal. Subsequently, the Positive and Negative Affect Schedule (37) (PANAS))was used to assess the participants’ current affective state. This was followed by task instructions for the upcoming tasks and a short training of those.

To further increase participants’ subjective hunger and ghrelin concentrations (38) in the no-meal session, they were asked to smell and to identify an odour probe (either a small amount of real peanut butter or banana, the latter of which was presented via a Sniffin’ Sticks pen (39)). Afterwards and approximately one hour after the first blood collection, the second blood sample was taken (T1).

Functional imaging was conducted after a 10-minutes indoor walk to Oslo University hospital. To keep participants’ stomach filled and ghrelin concentrations constant in the liquid-meal session, participants ate a banana shortly before scanning started. Participants completed several training trials of the first two fMRI tasks to become familiar with the response devices. After scanning, a third blood sample was collected (T2).

### 2.3 Ghrelin analysis

To measure plasma ghrelin concentrations, 2 ml EDTA tubes were prepared with 100μl protease inhibitor (Pefabloc^®^ SC Plus, Merck KGaA, Germany) before blood collection. All blood samples were put on ice immediately after venepuncture, then centrifuged for 15 minutes at 4°C and 3200g. Samples were then aliquoted and ghrelin plasma was stabilized with HCl before storage at −80°C in in-house freezer facilities. Ghrelin samples were analysed at the Hormone Laboratory at Oslo University Hospital (Oslo, Norway). Active (acylated) ghrelin concentrations were determined using the EZGRA-88K kit (Merck, Germany) in duplicates (total analytical CV at 488 pg/ml 12%).

### 2.4 Tasks

#### 2.4.1 Social recognition by experts (task 1)

This task, which we call “social-recognition-by-experts” task, was adapted from (40–42). In this task, participants are receiving two types of reward based on their personal music preferences: social reward and reward provided by an arbitrary computer algorithm. Social reward in the form of social recognition is present when the preferences of two music experts correspond to, and thereby endorse, participants’ music choices. Non-social or computer reward is present when a computer algorithm picks the same song as participants had previously chosen and assigns a virtual token to it. These tokens awarded by the computer were transformed into a physical reward at the end of the study (a USB stick with participants’ favourite songs in high audio resolution to take home). Hence, the receipt of the USB stick was only linked to the computer reward outcomes.

A few days before the first test session participants had sent in a list of 40 of their favourite music songs to the experimental team in order to have these songs evaluated by two supposedly renowned music experts. Additionally, participants had sent a picture of themselves, which was then converted to grayscale and cropped to be presented in the task.

Participants rated how much they liked a subset of 16 of their provided songs on a Likert-scale, ranging from “1=I do not like it” to “ 10=I love it” before each scanning session started. During task instructions prior to scanning, participants were shown photos and a description of the two music experts (a man called Ketil and a women called Sigrid) who ostensibly provided the expert ratings of the submitted music pieces (a translation from the original material in English into Norwegian is provided in Supplementary Materials). The descriptions provided in-depth information about the degree of expertise of each music expert across a range of popular music tastes. Afterwards, participants were required to rate how much each expert could be trusted to pick music they would like (“How much would you trust this person to pick music that you would like?”), and how much they would appreciate knowing that each expert liked listening to the same kind of music (“How much would you like it if you found out this person enjoyed the same music as you?), ranging from “ 1=not at all” to “7=very much”. Lastly, participants were informed that the experts had listened to all the songs and had provided reviews for each song. Reviews were preferences between each of the participant-provided songs and an alternative song, provided by the experimental team (these songs were chosen from Austrian and German artists who were presumably not known by an average Norwegian audience; two parallel versions of these experimenter-provided songs that were presented in a counterbalanced order in liquid-meal and no-meal sessions were available). Each participant-provided song was reviewed six times by the experts, relative to randomly selected alternative songs. As in the original studies, it was checked before the scanning session whether all the alternative songs were unknown to the participants. If not, the respective song was replaced by another one. This procedure was implemented to increase the likelihood that participants would choose their own songs over the alternative unknown songs when conducting the task in the scanner.

The social-recognition-by-experts task (task 1) was administered as the first of four experimental tasks in the scanner. It consisted of two runs with 48 trials each. At the beginning of each trial, participants were presented with two song titles at the top of the screen: one was selected from the participant’s list, the other one was selected from the experimenters’ list (see Figure 1). The two song titles were randomly presented on the left and right side of the screen. Pictures of the two experts were displayed in the middle of the screen and below each other. The participant’s picture was presented at the bottom of the screen, underneath the expert pictures. The words “I prefer” in Norwegian were placed under each picture. Participants were instructed to move their own picture beneath the song they liked best and wanted to own in “hi-fi” audio quality by pressing pre-assigned buttons on an fMRI response device placed in their right hands (ResponseGrips, Nordic Neurolab, Norway). Participants were given 3.6 sec to make a choice, otherwise a warning message was displayed asking for a faster response, and the next trial started after a 2 sec fixation cross. Within the 3.6 sec, participants’ pictures were moved underneath the chosen song title after their button press and a black-and-white scrambled picture was placed underneath the non-chosen song – this phase was termed “decision phase”. Next, the “social reward outcome phase” followed in which the pictures of each expert were moved underneath one of the song titles as an indication of their choice (with scrambled pictures being placed underneath the non-chosen songs).

There were three possible scenarios after the participants’ and experts’ choices were presented (given that participants had chosen “their” songs): (1) Both experts preferred the participant-provided song – reviewers’ agreement with the current choice signifying “social reward”. (2) Both experts preferred the alternative song – reviewers’ disagreement with the current choice signifying the omission of social reward. (3) The experts showed diverging preferences, one had chosen the participant-provided song while the other one had not. In each run, 20 trials resulted in reviewers’ agreement, 20 trials in reviewers’ disagreement, and 8 trials in diverging reviewers’ preferences. The social reward outcome phase lasted for 2 sec.

Next, the song titles alternately changed color between green and white (every 50ms, for 1s) before a song was chosen by a computer algorithm. Its title was presented at the bottom of the screen, thereby assigning a token to said song (duration 2 sec). This phase was termed the “computer outcome phase”. Participants were told that songs they chose had a slightly higher chance of receiving a token from the computer algorithm (51%) at the end of the trial to motivate them to choose according to their real music preferences. They were also informed that the songs with the most tokens would be given to them in a digital high-quality version at the end of the experiment to make the virtual token motivationally relevant. In reality, each song had a 50% chance of being selected by the computer, meaning that half of the trials resulted in a win (the computer selected the participant-provided song), and the other half of the trials in no-win (the computer selected the experimenter-provided song). The trials were separated by an inter-trial interval in which a white fixation cross was presented centrally for 2 sec. The task was programmed and presented in E-Prime 2.0/3.0 (Psychology Software Tools, Inc., Sharpsburg, PA). Each run took about 9 min. Figure 2 demonstrates the timing of one trial.

**Figure 2.**
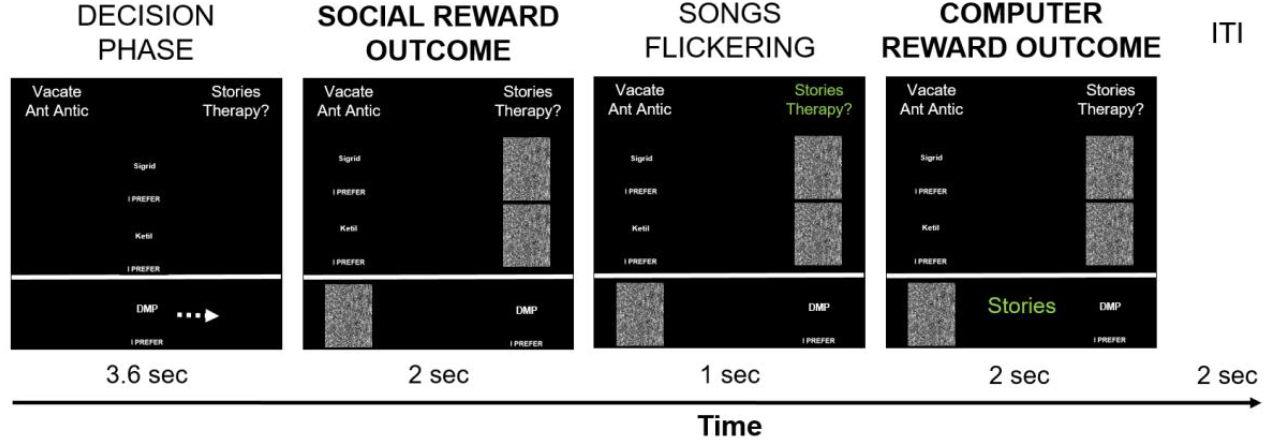
Timing of one social-recognition-by-experts-trial. Each trial started with the presentation of two song titles on top of the screen, of which one was provided by the participant, and one was provided by the experimenters. Below, pictures of the two supposed music experts (replaced here with the placeholder characteres “Sigrid” and “Ketil” and the participant (placeholder characters for author DMP) were shown in the middle of the screen. Participants had 3.6 sec to choose which song they liked best and would like to own in hi-fi data quality by pressing pre-defined buttons on the response device to move their picture either to the left or to the right, under their preferred song. Participants’ choice is implied here with a white arrow (not present during scanning), indicating that the participant-provided song was chosen – signifying the decision phase. Afterwards, the choices of the music experts are shown by moving their pictures under their preferred song – the social reward outcome phase. In the current example, both experts chose the song provided by the experimenters, thereby signifying reviewers’ disagreement with the participant’s preference in this trial. Subsequently, the names of the two songs alternately changed colour ten times from white to green, for 50 ms for each colour, signifying the choice period of the computer. After this flickering phase, the song chosen by the computer algorithm was presented on the bottom of the screen in green colour – the computer reward outcome phase. In the current example, the participant-provided song was chosen by the computer algorithm and a token was assigned to said song, indicating a win. The computer reward outcome phase always followed the social reward outcome phase, but their respective outcomes were independent from each other. Trials were separated by a 2 sec inter-trial-interval with a white fixation cross presented centrally.

After the end of scanning, participants were again asked to rate the 16 respective songs of the current session (outside the scanner) in terms of how much they liked them. This was done to assess whether participants had changed their mind in response to the experts’ reviews. Afterwards, participants provided further ratings on how much they trusted the experts’ choices to be similar to their own (“How much would you trust this person to pick music that you would like?”), and how much they would like it if the experts had a similar taste in music (“How much would you like it if you found out this person enjoyed the same music as you?”), again rated from “1=not at all” to “7=very much”. A full list of all rating questions is provided in Supplementary Materials. At the end of each test session, participants were given a USB stick with high-quality digital versions of five of their provided songs, which were purchased for them.

#### 2.4.2 Social affirmation task (task 2)

The social affirmation task was designed to assess behavioural and neural dimensions of social reward. It was the third task presented in the scanner and started approximately 35 minutes after task 1.

Participants were told that they were going to read several statements on the screen and then be asked to rate the pleasantness they experienced while reading the statement. Instructions did not mention that all statements referred to the participants or that the statements were given by the experimenters.

Each trial started with the jittered presentation of a fixation cross (duration between 12.25 and 14 sec minus the response time from the previous trial; uniform distribution of 250 ms intervals) – see Figure 3. Each statement was presented centrally on the screen for 3 sec. Afterwards, participants answered the question “How did you experience the statement?” on a visual analogue scale (VAS) with the end points “unpleasant” (coded as 0) and “pleasant” (coded as 100). Participants were given 12 sec to respond; otherwise, they were reminded to respond faster in the next trial. Responses were given by pressing pre-assigned buttons on fMRI response devices placed in both hands. The task consisted of 16 neutral statements (neutral condition) and 16 compliments (social reward condition) which were presented in a randomized order.

**Figure 3.**
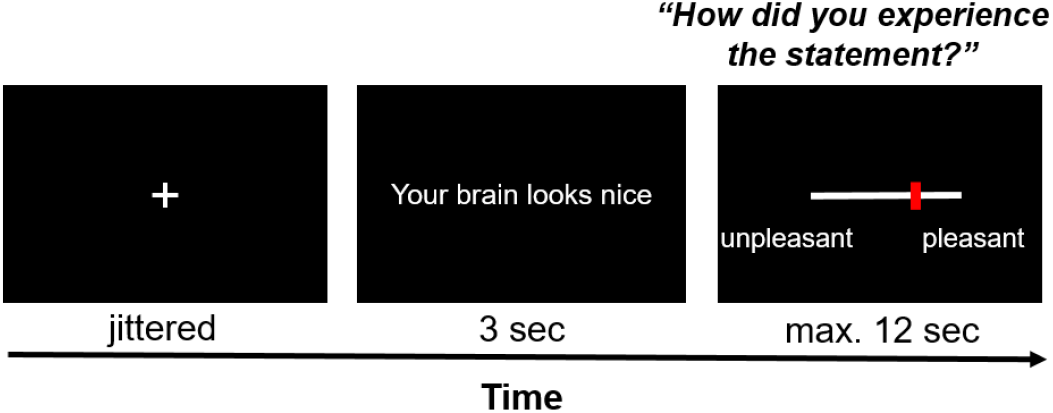
Exemplary trial of the social affirmation task. After a jittered inter-stimulus interval, a statement was presented on the screen for 3 sec, the pleasantness of which participants rated on a VAS scale with *ResponseGrips* (Nordic Neurolab, Norway).

A list of the presented neutral statements and compliments can be found in Supplementary Materials. The neutral statements described either physical (“Your brain consists of 73% water”) or behavioural features (“You are a participant in an experiment”) of the participant. Similarly, the compliments referred to physical (“Your head has the perfect form to fit into the scanner coil”) or behavioural features (“You are an exemplary participant”). The word length and structure of the neutral statements and compliments was kept as parallel as possible to guarantee that both only differed in terms of whether they provided social affirmation or not.

Two parallel versions of the task were created and presented in a pseudo-randomised order in the liquid-meal and the no-meal session. Task duration was about 9 min.

Internal consistency of the pleasant ratings was very high with alpha= 0.958 for the compliments items, and alpha=0.947 for the neutral statements. Factor analysis with direct oblimin rotation was performed separately on the compliments and the neutral statements. Scree plots indicated a clear one-factor-solution per scale, which explained 62.332% of the variance in the ratings of compliments, and 57.786% of the variance in the ratings of neutral statements.

### 2.5 Additional measures

Participants also completed a set of self-reported measures online before the first test session. A list of these questionnaires can be found in the preregistration. Their results will be presented elsewhere.

### 2.6 fMRI data acquisition and preprocessing

Imaging was conducted on a Philips Ingenia 3 T MRI scanner (Philips Healthcare, Best, NL). For functional imaging, we used a 32-channel SENSE head-coil using an echo planar imaging (EPI) sequence with the following parameters: voxel size = 3 mm, repetition time (TR) = 2500 ms, echo time (TE) = 30 ms, flip angle = 90°, FOV =240×240×120, 40 slices, interleaved without gap. In task 1, 210 functional volumes were acquired per run; in task 2, 212 volumes were acquired once. At the beginning of each task, five dummy volumes were scanned. For the high-resolution anatomical image, a T1-weighted 3D MP-RAGE sequence was used with the following parameters: TR/TE = 5.2/2.3 ms, flip angle = 8°, voxel size = 1 x 1 x 1 mm, FOV = 184×256×256; scan duration: approx. 5 min.

For all the neuroimaging analysis, including preprocessing, the software Statistical Parametric Mapping (SPM12, Wellcome Trust Centre for Neuroimaging,https://www.fil.ion.ucl.ac.uk/spm/software/spm12/) running on MATLAB Version R2019a was used.

fMRI pre-processing and first level analyses were performed on the high-performance computing resource Saga, owned by the University of Oslo, and operated by the Department for Research Computing at USIT, the University of Oslo IT-department (http://ww.hpc.uio.no/).

Data pre-processing was carried out separately for each scanning session and task. Original data were converted from dicom to nifti format with MRIcroGL 1.2.20190902++ (43), individual brain anatomy files were anonymized using the De-face image function that deletes image information of facial features. This was required by the local ethics board at a later stage and thus not included in the preregistration. Other preprocessing steps included slice time correction (to the first slice), motion correction (referenced to the mean image) and unwarping, spatial normalization to MNI (Montreal Neurological Institute) stereotactic space, and spatial smoothing (6 mm FWHM Gaussian kernel). Default algorithms and parameters were used for preprocessing. We applied a 4 mm threshold for excessive head movements, which none of the included participants in the final samples exceeded. The ArtRepair toolbox (https://cibsr.stanford.edu/tools/human-brain-project/artrepair-software.html) was only used to identify scans affected by artefacts caused by an unexpected interaction between the current EPI sequence and a movement-correction setting used during everyday clinical routine measurements at Oslo University Hospital. The data from several participants showed non-correctable signal distortions in either the liquid-meal or the no-meal sessions. In the recognition-by-experts task, twelve participants were affected by these artefacts, leaving us with 48 complete datasets for neuroimaging analyses. In the social affirmation task, nine participants were affected, leaving us with 52 complete datasets for neuroimaging analyses.

### 2.7 Analysis

All the behavioral and hormonal analyses were conducted with SPSS 27, Jamovi 1.6.23 (70) and JASP 0.16.3 (71). The reported analysis and results follow the preregistration and any deviations from it are clearly stated. Null effects of ghrelin were followed up with Bayesian correlation analyses (via non-parametric Kendall’s tau) to provide evidence for the probability of these null effects. The Bayesian analyses were not preregistered. All data, code, and materials will be made publicly available via the Open Science Framework after acceptance.

#### 2.7.1 Manipulation check

To verify the effects of liquid meal intake, we analysed variation in plasma ghrelin concentrations and self-report ratings as indices of nutritional state changes due to the meal. Deviating from the pre-registration, we conducted a linear mixed model to analyse ghrelin concentrations instead of a repeated-measures ANOVA because linear mixed models model individual data more accurately and can also include participants with missing data (44,45). This is relevant in the current study as plasma ghrelin samples could not be collected in all participants at all measurement time points due to complications when collecting the blood. Ghrelin concentrations were analysed with a linear mixed model with nutritional state (liquid-meal/no-meal) and the three sample time points (T0/baseline, T1, T2) as fixed factors. The random effects structure included a random intercept for participant and a random slope for nutritional state (model description: ghrelin concentrations ~ 1 + nutritional_state + time_points + nutritional_state:time_points + (1 + nutritional_state | participant). The Satterthwaite method for approximation of degrees of freedom was used and a restricted maximum likelihood estimation for fixed effects was applied. Post-hoc tests were Bonferroni-corrected. As effect size measures, we report semi-partial R^2^ (46), for which values of 0.02, 0.13, and 0.26 denote small, medium and large effects (47).

For self-report ratings of current bodily states, we first computed a composite score of all questions and compared it for liquid meal vs. no-meal. We then conducted this comparison for each of the questions asked separately, either with paired t-tests or with non-parametric alternatives in case the residuals were not normally distributed. The same strategy was applied to affective state ratings (PANAS) and the remaining self-report ratings.

#### 2.6.2 Task 1: behavioral analysis

Following (40), we calculated an index of how much the reviewers’ opinions influenced participants’ subjective song liking, i.e. a measure of susceptibility towards the reviewers’ preferences. This index was denoted as *B_inf_* and was calculated for each participant separately for liquid-meal and no-meal. Binf reflects the change (denoted in the standard deviations) by which the individual liking ratings of each song increased or decreased after the experiment in relation to reviewers’ opinions of the respective song (see (41) for a detailed description). The more positive the value of Binf, the more participants changed their opinion according to the reviewers’ preferences. Binf indexes were compared for liquid-meal and no-meal with a paired t-test. Only 48 participants were available for this analysis because we did not collect the liking ratings in the first participants, or the administration either before or after the scanning session was not conducted.

Repeated-measures ANOVAs were used to analyse the ratings on trust and appreciation towards the reviewers. Nutritional state (liquid-meal vs. no-meal), time of assessment (pre- and post-task) and reviewer identity (Ketil vs. Sigrid) served as within-subject factors. For these calculations, data of 60 participants were available.

#### 2.7.4 Task 1: neuroimaging analysis

Following the preprocessing, first-level analysis of the data of each participant was performed based on the General Linear Model framework as implemented in SPM12 (48). From the 96 trials in the recognition-by-experts task, only those trials were included in further analyses in which participants had chosen their preferred song. In the social outcome phase, 40 trials could reflect reviewers’ agreement (“social reward”), while 40 trials could reflect reviewers’ disagreement, and 16 trials could reflect diverging reviewers’ preferences given that participants had chosen “their” songs. Of note, the 16 trials with diverging reviewers’ preferences were not included in subsequent analyses. In the computer outcome phase, 48 trial could provide a win (when the participant’s song was chosen), while 48 trials could provide a no-win. Trials in which participants were too slow to make a choice were not usable for further analysis (1.1% of all trials in liquid-meal sessions, 1.5%of all trials in no-meal sessions). Reaction times did not differ between liquid-meal and no-meal sessions (p=.728; liquid-meal: mean=1323.84 ms, SD=212.45; no-meal: mean=1331.89 ms, SD=229.86).

Seven regressors of interest were convolved with SPM’s canonical hemodynamic response function with a duration of 2 sec. Five of them corresponded to the main task conditions: reviewers’ agreement, reviewers’ disagreement, reviewers’ diverging preferences in the social reward outcome phase; and win and no-win in the computer outcome phase. Two further regressors were included to take care of those trials in which participants chose the alternative song rather than the one they had provided themselves (one in the social reward outcome phase, one in the computer outcome phase). One nuisance regressor was included to model the last scanning volumes in cases where participants had missed responses, which shortened the E-Prime paradigm but not the scanning time. In addition to those regressors, six nuisance regressors representing the realignment parameters were also included in the first-level model to account for residual motion artefacts. The reported first-level analysis is different from what was pre-registered because we decided to model the events of the social and the computer outcome phase separately (corresponding to the analysis reported in (42). This is in contrast to the original study by (40) which was focused on social influence on non-social reward and thereby modelled all six possible combinations of social and computer outcomes during the computer reward outcome phase, as well as a nuisance regressor for the decision phase of the participant. After model estimation, reviewers’ agreement, reviewers’ disagreement, win and no-win were modelled against the implicit baseline, as well as the following contrasts of interest: reviewers’ agreement > reviewers’ disagreement, win > no-win.

Group level analysis consisted of three steps. First, we assessed whether the previous findings could be replicated (40–42). To this aim, we computed two one-sample t-tests, one with the social reward contrast (agreement > disagreement), the second one with the computer reward contrast (win > no-win). Only data from the liquid-meal sessions were considered for this replication analysis because this nutritional state can be considered the default state in which participants in the original studies were tested. To verify previously reported activation in reward-related areas in the recognition-by-experts task (40–42), we used small volume correction (SVC) after building spheres with an 8 mm radius centered on the previously reported peak MNI coordinates. The regions were generated using Marsbar (49). The complete list of regions is reported in Table 1 and included: bilateral ventral striatum (VS), ventromedial prefrontal cortex (vmPFC), and bilateral lateral orbitofrontal cortex (lOFC). Starting threshold was set at p < .001. Small volume correction was set to p<.05 FWE, peak-level. This replication analysis was not pre-registered, but we believe it to be a fundamental step to first verify that the task elicited activation in reward-related brain regions reported in the previous studies before analyzing whether brain activity in these regions varies depending on ghrelin variation and nutritional state.

**Table 1.** List of MNI coordinates used to generate spheres with 8 mm radius for SVC analysis. The studies from which they were taken and the phases for which they were reported significant are also listed. *Regions showing significant activity and thus subsequently employed for ROIs analysis to test the main hypotheses of whether liquid-meal vs. no-meal and ghrelin variation had an impact on brain activity in the reward network.

| ROIs | x | y | z | Study | Phase |
| --- | --- | --- | --- | --- | --- |
| **Left Ventral striatum\*** | -10 | 8 | -12 | Campbell-Meiklejohn (40) | Social reward outcome |
| **Right Ventral striatum** | 8 | 8 | -12 | Campbell-Meiklejohn (40) | Social reward outcome |
| **Left Ventral striatum** | -16 | 16 | 2 | Campbell-Meiklejohn (40) | Computer reward outcome |
| **Right Ventral striatum\*** | 14 | 10 | -8 | Campbell-Meiklejohn (40) | Computer reward outcome |
| **vmPFC\*** | 0 | 58 | -6 | Tobler (42) (taken from 50) | Social & computer reward outcome |
| **Left IOFC** | -33 | 28 | -16 | Campbell-Meiklejohn (41) | Social & computer reward outcome |
| **Right IOFC** | 36 | 33 | -10 | Campbell-Meiklejohn (41) | Social & computer reward outcome |

As a second step, we compared brain activation in the liquid-meal and no-meal sessions for the social reward outcome phase (agreement > disagreement) and for the computer reward outcome phase (win > no-win) on the whole-brain level with two paired-samples t-tests. Starting threshold was set at p < .001 and significant threshold was set at p<.05, FWE corrected, peak-level.

As a third step of the group-level analyses, we performed a regions-of-interest (ROIs) analysis where we focused on reward-related areas that showed significant activity in the replication analysis (described above; vmPFC, bilateral VS). For each ROI, we extracted the mean activity for each participant in the liquid-meal and the no-meal session from the baseline contrasts for reviewers’ agreement, reviewers’ disagreement, win, and no-win. Three-way repeated-measures ANOVAs with the within-subject factors nutritional state (liquid-meal, no-meal), reward outcome category (social reward outcome, computer reward outcome), and outcome valence (favourable outcome, unfavourable outcome) were computed for each ROI. Multiple comparison corrections were based on the number of ROIs involved in the analyses (Bonferroni-corrected p < .017).

#### 2.7.5 Task 1: Relationship between ghrelin concentrations at T1, susceptibility to social recognition and brain activation during social and computer reward outcomes

In order to investigate the relationship among behavioral, hormonal, and brain processes, we conducted Spearman correlations. All correlations were conducted on the difference values between liquid-meal and no-meal sessions to capture individual variation of these differences. This is denoted with the symbol “Δ” in front of each measure in the results section. The boxplot function in SPSS was used to identify outlier values (beyond the

1.5 interquartile range) in each variable. In contrast to the preregistration, we removed these outliers from subsequent correlation analyses instead of applying a winsorisation procedure because they were rather extensive. We correlated the susceptibility index Binf with plasma ghrelin concentrations at T1 to test whether the ghrelin satiation response (i.e., the difference between ghrelin concentration following meal intake and ghrelin concentration at a similar time point when participants were continuously fasting) was associated with how much the participants changed their song liking due to the reviewers’ preferences in the two sessions. Further correlations were calculated between the ghrelin satiation response and the difference in mean ROI brain activity for the two sessions. Lastly, correlations were calculated between the susceptibility index Binf and ROIs’ brain activity. Multiple comparison corrections were based on the number of ROIs involved in the analyses, separately in the social and computer outcome phases (three ROIs per analysis resulted in a corrected p-value of p < .017).

#### 2.7.5 Task 2: behavioral analysis

To test the effect of ghrelin concentrations and nutritional state on the pleasantness of the statements, a two-steps approach was adopted (deviating from the pre-registration, we did not model nutritional state and ghrelin concentrations in the same model because the two variables share variance with each other). First, we used nutritional state as an independent variable to predict pleasantness in a linear-mixed model (LMM) analysis. Specifically, single-trial pleasantness ratings were modelled as a function of statement type (compliments/neutral statements), nutritional state (liquid-meal/no-meal), the mean-centered trial number, and the interaction of statement type x nutritional state as fixed effects, including a random intercept for participant and random slopes for statement type and nutritional state (model description: ratings ~ 1 + statement_type + nutritional_state + trial_number + statement_type:nutritional_state + (1 + statement_type +nutritional_state | participant)). Second, we investigated the effect of ghrelin concentrations on pleasantness ratings by computing a linear mixed model in which single-trial pleasantness ratings were modelled as a function of statement type (compliments/neutral statements), measurement session (1,2), mean-centered trial number, and mean-centered ghrelin concentrations (sample time point T1), and the interaction of statement type x ghrelin as fixed effects. This model included a random intercept for participant and random slopes for statement type and measurement session (model description: ratings ~1 + statement_type + measurement_session + trial_number + ghrelin + statement_type:ghrelin + (1 + statement_type + measurement_session | participant)). For both analyses, 60 participants were available with both sessions and six participants with one session (4 liquid-meal and 2 no-meal).

#### 2.7.6 Task 2: neuroimaging analysis

First-level analysis for task 2 was again performed based on the General Linear Model framework as implemented in SPM12. Three regressors were computed to model the onset of trial events: one for compliments, one for neutral statements (both with a duration of 3 sec), and one for the pleasantness ratings (trial-wise response times as duration). A nuisance regressor modelled the last five volumes at the end of the task. In addition to those regressors, six nuisance regressors representing the realignment parameters were also included in the model to account for residual motion artefacts. After model estimation, compliments and neutral statements were modelled against the implicit baseline and two contrasts were created, as well as the contrast compliments > neutral statements.

For the group-level analysis, we first wanted to investigate whether social affirmation elicits activation in reward-related brain areas on the whole brain level. Therefore, we computed a one-sample t-test for the contrast compliments > neutral statements in the liquid-meal session only, as this should be considered the default nutritional state in typical experimental set-ups.

As a second step, we compared the compliments > neutral statements contrast for liquid-meal and no-meal sessions on the whole-brain level. Here, a paired-sample t-test was performed. For both analyses, whole-brain results are reported at p < .05 FWE, peak-level. As a last step of the group-level analyses, a ROIs analysis was performed using the same ROIs as the ones used in the analysis of task 1 (limited to the social reward outcome phase: bilateral VS and vmPFC). This will allow for better comparability between the results of task 1 and 2. Mean activity within each ROI was extracted for each participant from the baseline contrasts for compliments and neutral statements, in the liquid-meal and the no-meal session. Repeated-measures ANOVAs with the within-subject factors nutritional state (liquid-meal, no-meal) and statement type (compliments, neutral statements) were calculated. Multiple comparison corrections were based on the number of ROIs involved in the analyses (Bonferroni-corrected p < .017).

#### 2.7.7 Task 2: Relationship between ghrelin concentrations at T1, pleasantness ratings and brain activation during compliments and neutral statements

Analogue to task 1, we tested the relationship among behavioural, hormonal, and neuroimaging measures with Spearman correlations. Again, correlations were conducted on the difference values between liquid-meal and no-meal sessions (denoted with the symbol “Δ”) and outlier values were removed from subsequent analyses after identification with the boxplot function. First, correlations between the ghrelin satiation response (i.e., the difference in ghrelin following meal intake vs. when continuously fasting) and pleasantness ratings were calculated, separately for compliments and neutral statements. Second, ROI brain activation was correlated with pleasantness ratings, again separately for compliments and neutral statements. Third, the ghrelin satiation response and ROI brain activation was correlated. Corrections for multiple comparisons were applied as in task 1 (three ROIs per analysis resulted in a Bonferroni-corrected p-value of p < .017).

## 3. Results

### 3.1 Manipulation check

The linear mixed model testing ghrelin variation during the experiment showed significant effects of nutritional state (b=82.1, SE= 21.0, t(59.0)=3.92, p<.001, semi-partial R^2^=0.21) and sample time point (T0/baseline vs. T1: b=-203.2, SE=21.0, t(227.1)=-9.67, p<.001, semi-partial R^2^=0.32; T1 vs. T2: b=69.3, SE=18.0, t(226.9)=3.84, p<.001, semi-partial R^2^=0.32). Moreover, the nutritional state x sample time point interaction was significant (both p-values <.001, both semi-partial R^2^=0.29). Bonferroni-corrected post-hoc tests demonstrated a significant ghrelin satiation response at T1, about 30 minutes after the liquid-meal. On average, plasma ghrelin concentrations were dampened by more than 300 pg/ml after participants had consumed the liquid-meal compared to the same time point in the no-meal session. In contrast, no differences between liquid-meal and no-meal in ghrelin concentrations were observed for T0/Baseline and T2 (both p-values>.999); see Table 2 and (27) for more details.

**Table 2.** Plasma concentrations of acylated ghrelin (pg/ml)

|  | Liquid-meal |  |  | No-meal |  |  |
| --- | --- | --- | --- | --- | --- | --- |
|  | *M* | *SD* | *n* | *M* | *SD* | *n* |
| *Baseline (T0)* | 906 | 518 | 62 | 851 | 483 | 60 |
| *T1* | 502 | 274 | 57 | 836 | 445 | 52 |
| *T2* | 843 | 467 | 61 | 779 | 403 | 56 |

Blood glucose concentrations measured at the beginning of each test session did not differ significantly from each other (t(59)=−1.86, p=.067, d=−0.24). All subjective reports on bodily states differed significantly between liquid-meal and no-meal (all p’s < .001). Time since last meal before the experiment did not differ between the two test sessions (p=.210; please note that ten participants were not included in this calculation as they either had misunderstood the question or had filled in impossible values); see Table 3 for details.

**Table 3.**
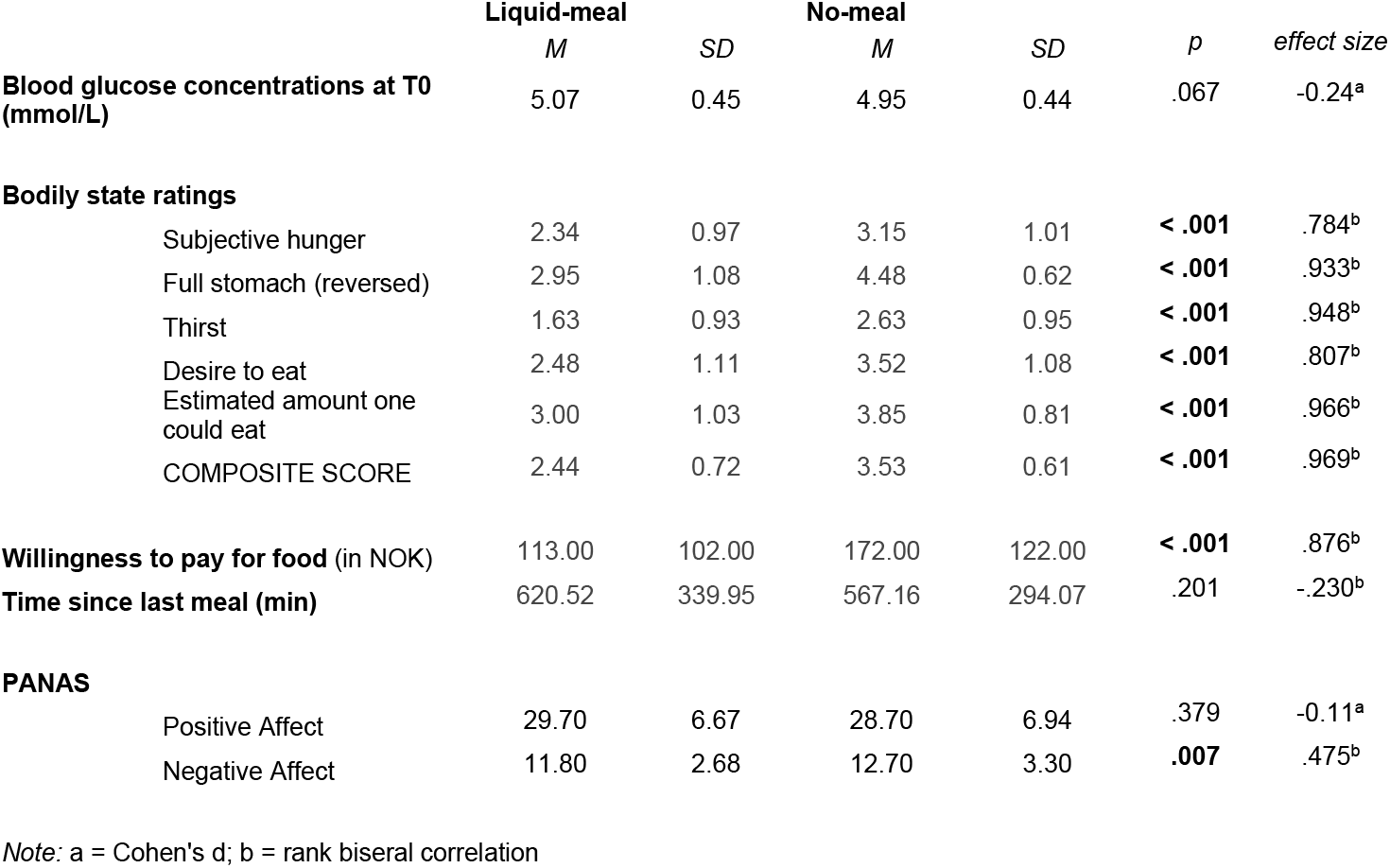
Effects of liquid-meal and no-meal on bodily and affective states.

|  | Liquid-meal |  | No-meal |  | *p* | effect size |
| --- | --- | --- | --- | --- | --- | --- |
|  | *M* | *SD* | *M* | *SD* |  |  |
| **Blood glucose concentrations at T0 (mmol/L)** | 5.07 | 0.45 | 4.95 | 0.44 | .067 | -0.24<sup>a</sup> |
| **Bodily state ratings** |  |  |  |  |  |  |
| Subjective hunger | 2.34 | 0.97 | 3.15 | 1.01 | < .001 | .784<sup>b</sup> |
| Full stomach (reversed) | 2.95 | 1.08 | 4.48 | 0.62 | < .001 | .933<sup>b</sup> |
| Thirst | 1.63 | 0.93 | 2.63 | 0.95 | < .001 | .948<sup>b</sup> |
| Desire to eat | 2.48 | 1.11 | 3.52 | 1.08 | < .001 | .807<sup>b</sup> |
| Estimated amount one could eat | 3.00 | 1.03 | 3.85 | 0.81 | < .001 | .966<sup>b</sup> |
| COMPOSITE SCORE | 2.44 | 0.72 | 3.53 | 0.61 | < .001 | .969<sup>b</sup> |
| **Willingness to pay for food (in NOK)** | 113.00 | 102.00 | 172.00 | 122.00 | < .001 | .876<sup>b</sup> |
| **Time since last meal (min)** | 620.52 | 339.95 | 567.16 | 294.07 | .201 | -.230<sup>b</sup> |
| **PANAS** |  |  |  |  |  |  |
| Positive Affect | 29.70 | 6.67 | 28.70 | 6.94 | .379 | -0.11<sup>a</sup> |
| Negative Affect | 11.80 | 2.68 | 12.70 | 3.30 | .007 | .475<sup>b</sup> |

Note: a = Cohen's d; b = rank biserial correlation

### 3.2 Task 1: behavioral analysis

When comparing the susceptibility index *B_inf_* between liquid-meal and no-meal, no significant difference was observed (t(46)=.985, p=.330).

Concerning the trust and appreciation ratings of the two experts, results showed a main effect of time of assessment for both questions, with a decrease from pre- to post-task (trust: F(1,59)=21.717, p<.001, ηp^2^=.269; appreciation: F(1,59)=13.786, p<.001, ηp^2^=.189). Nutritional state had no effect on these ratings (all p’s > .136), and neither did expert identity (all p’s > .056).

### 3.3 Task 1: neuroimaging results

Previous findings were partly replicated for both the social reward outcome contrast and the computer reward outcome contrast in the liquid-meal session. For the social reward outcome, we found significantly higher activity, upon small volume correction, in the left ventral striatum (p<.001, MNI coordinates: −8, 8, −8) and in the vmPFC (p=.009, MNI coordinates: −4, 62, −8) for reviewers’ agreement compared to their disagreement. For the computer reward outcome, we found significantly higher activity in the right ventral striatum (p=.030, MNI coordinates: 12, 14, −6) and in the vmPFC (p=.005, MNI coordinates: 0, 56, 0) for win compared to no-win.^1^

No significant differences were detected in the whole-brain analysis comparing liquid-meal and no-meal sessions, for both the social and the computer reward outcomes.

The subsequent ROIs analyses (full-factorial repeated measures ANOVAs with factors nutritional state, reward outcome category, and outcome valence) were calculated with extracted time series of regions (vmPFC, right and left VS) showing significant activation differences in the replication analysis (Bonferroni-corrected p < .017). The vmPFC ANOVA did not show any significant result as the main effect of outcome valence did not pass significance (F(1,47)=5.82, p=.020, η_p_^2^=0.11). Descriptively, vmPFC activation was higher during favourable than unfavourable outcomes. All other main effects or interactions were not significant either (all p’s > .092). The right VS ANOVA showed a significant main effect of outcome valence in the right VS ROI (F(1,47)=15.67, p<.001, η_p_^2^=0.25) with higher activation during favourable than unfavourable outcomes (all other p’s > .100). In the left VS ANOVA, significant main effects were observed for outcome valence (F(1,47)=30.68, p<.001, η_p_^2^=0.40) and reward outcome category (F(1,47)=7.92, p=.007, η_p_^2^=0.14). Left VS activation was higher for favourable than unfavourable outcomes, and for computer reward than social reward outcomes (all other p’s > .038, see Table 4 and Figure 4).

**Table 4.** Means and SD of extracted time series ROI activation

| **TASK 1 (n=48)** |  | **vmPFC** |  | **R VS** |  | **L VS** |  |
| --- | --- | --- | --- | --- | --- | --- | --- |
|  |  | *M* | *SD* | *M* | *SD* | *M* | *SD* |
| *liquid-meal* | *social - agreement* | -0.29 | 1.36 | -0.19 | 0.72 | -0.25 | 0.54 |
|  | *social - disagreement* | -0.30 | 1.24 | -0.34 | 0.69 | -0.53 | 0.64 |
|  | *non-social - win* | -0.18 | 0.57 | -0.04 | 0.33 | -0.09 | 0.29 |
|  | *non-social - no-win* | -0.37 | 0.55 | -0.11 | 0.40 | -0.21 | 0.31 |
| *no-meal* | *social - agreement* | 0.06 | 0.92 | -0.04 | 0.64 | -0.20 | 0.60 |
|  | *social - disagreement* | -0.07 | 0.97 | -0.16 | 0.67 | -0.36 | 0.51 |
|  | *non-social - win* | -0.11 | 0.63 | -0.04 | 0.30 | -0.13 | 0.31 |
|  | *non-social - no-win* | -0.26 | 0.64 | -0.07 | 0.31 | -0.22 | 0.32 |
| **TASK 2 (n=52)** |  | **vmPFC** |  | **R VS** |  | **L VS** |  |
|  |  | *M* | *SD* | *M* | *SD* | *M* | *SD* |
| *liquid-meal* | *compliments* | 0.12 | 0.54 | -0.05 | 0.32 | -0.09 | 0.32 |
|  | *neutral statements* | -0.15 | 0.62 | -0.02 | 0.23 | -0.06 | 0.26 |
| *no-meal* | *compliments* | 0.14 | 0.49 | 0.11 | 0.31 | 0.06 | 0.33 |
|  | *neutral statements* | -0.21 | 0.56 | 0.03 | 0.33 | -0.03 | 0.36 |

**Figure 4.**
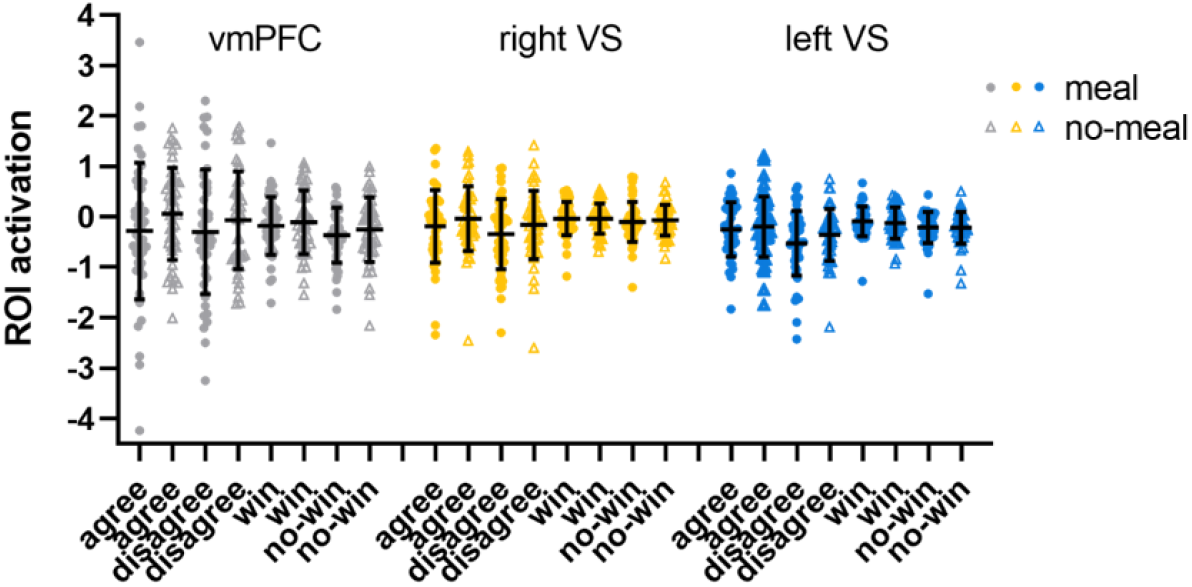
ROI activation in the recognition-by-experts task. Black horizontal bars represent mean values, error bars represent 1 standard deviation. Single-subject data are plotted as dots (liquid-meal condition) or triangles (no-meal condition) for social reward (agree, disagree) and computer reward (win, no-win). Please note that outlier values are included in this figure.

### 3.4 Task 1: Relationship between ghrelin concentrations at T1, susceptibility to social recognition, and brain activation during social and computer reward outcomes

Spearman correlations revealed a significant positive correlation between Δghrelin at T1 and ΔvmPFC activity in the computer reward outcome phase (win > no-win) (rs(37)=.438, p=.007; see Fig. 5), while no association was found between Δghrelin at T1 and ΔvmPFC activity in the social reward outcome phase (agreement > disagreement) (r_s_(36)=.002, p=.992). The larger the ghrelin suppression by the liquid-meal compared to no-meal for non-social computer reward, the larger the vmPFC suppression in the liquid-meal compared to the no-meal session. In contrast, participants with small or absent ghrelin suppression by the meal showed enhanced vmPFC activation in the liquid-meal compared to the no-meal session for non-social computer reward. None of the other correlations were significant (all p-values > .200).

**Figure 5:**
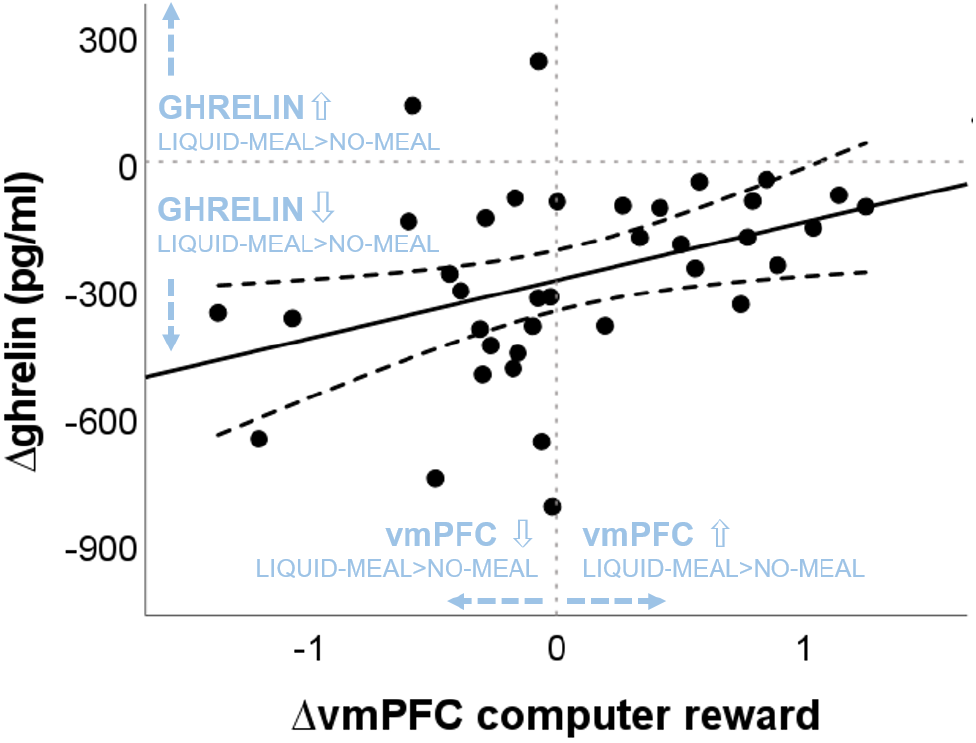
Association between nutritional-state dependent differences in vmPFC activation during computer reward outcomes and ghrelin concentrations. “liquid-meal minus no-meal” session differences of vmPFC activation during computer reward is plotted on the x-axis. Positive values indicate higher vmPFC activation in the liquid-meal than the no-meal session, while negative values indicate lower vmPFC activation in the liquid-meal than the no-meal session. On the y-axis, “liquid-meal minus no-meal” session differences in ghrelin concentrations at T1 are plotted (Δghrelin (pg/ml). Negative values indicate the size of the satiety response at T1 (i.e., the decrease in ghrelin concentrations after having a meal compared to no meal), while positive values indicate higher ghrelin concentrations at T1 in the liquid-meal than the no-meal session. The solid line depicts the regression line of the correlation, the dashed lines depict the 95% CI of the regression line.

Bayesian analyses showed moderate evidence for no association between Δghrelin and *ΔB_inf_,* and between Δghrelin and ΔvmPFC/ΔleftVS activation during social reward and between Δghrelin and ΔleftVS activation during computer reward outcomes, respectively (all BF01>3). Evidence for no association between Δghrelin and ΔrightVS activation during social reward and computer reward outcomes was only anecdotal. In contrast, evidence in favour of the observed positive association between Δghrelin and ΔvmPFC activity during computer reward outcomes was moderate with a BF_10_=5.858 (equivalent to BF_01_=0.171 in Table 5, which is calculated as 1/BF_10_: 1/5.858), see Table 5.

**Table 5.**
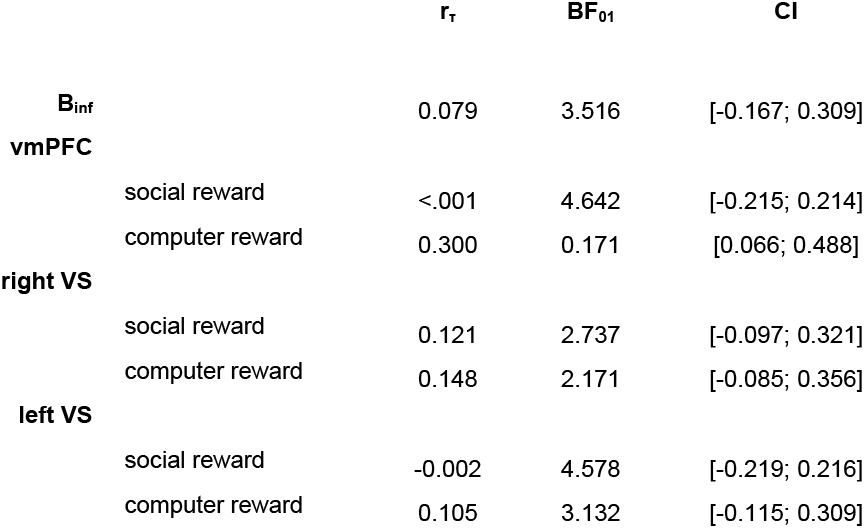

### 3.5 Task 2: behavioral results

Both models testing the effects of (1) nutritional state and (2) ghrelin concentrations on subjective pleasantness ratings showed that participants rated the compliments more pleasant than neutral statements, by more than 13 points on the rating scale (main effect of statement type in the nutritional state model: b=-13.99, SE=1.22, t(64.7)=-11.51, p<.001, semi-partial R^2^=0.67; main effect of statement type in the ghrelin model: b=−13.61, SE=1.23, t(59.6)=−10.49, p<.001, semi-partial R^2^=0.65). Both models also showed that experienced pleasantness decreased with increasing trial numbers (main effect of trial number in the nutritional state model: b=−0.09, SE=0.02, t(3832.2)=−4.81, p<.001, semi-partial R^2^=0.006; main effect of trial number in the ghrelin model: b=-0.09, SE=0.02, t(3280.6)=-4.68, p<.001, semi-partial R^2^=0.007). Neither a significant main effect of nutritional state, nor an interaction effect of nutritional state and statement type on pleasantness was found (both p-values > .131). In the ghrelin model, a main effect of measurement session was observed (b=-2.79, SE=0.84, t(46.1)=-3.34, p=.002, semi-partial R^2^=0.19). Pleasantness ratings were in general lower during the second test session by 2-3 points on the scale. Ghrelin concentration had no significant effect on pleasantness rating (b=-0.003, SE=0.002, t(76.3)=-1.67, p=.099). Bayesian analysis showed moderate evidence for no association between mean ghrelin concentrations at T1 and mean pleasantness ratings in a correlation analysis (BF_01_=5.858, r_τ_=0.020, CI = [-0.147; 0.184].

### 3.6 Task 2: neuroimaging results

First, we tested whether compliments elicited higher brain activation than neutral statements following the liquid meal to demonstrate the validity of the social affirmation task (by applying a whole brain analysis approach). This one-sample t-test showed a significantly higher activation in the medial prefrontal cortex (mPFC, MNI coordinates: 6, 54, 20) (p=.023 peak-level) for compliments than neutral statements. No other significant activation differences were observed. Second, testing for effects of nutritional state in the contrast compliments > neutral statements (paired-samples t-test, whole brain analysis), no significant brain activation was observed after multiple comparison correction.

Third, ROI analyses (full-factorial repeated measures ANOVAs with factors nutritional state and statement type) were conduced (Bonferroni-corrected p < .017). We observed a main effect of nutritional state in the right ventral striatum (F(1,51)=6.69, p=.013, η_p_^2^=.12), while this main effect did not pass significance in the left ventral striatum (F(1,51)=3.93, p=.053, η_p_^2^=.071). Descriptively, brain activity was lower in bilateral ventral striatum following the meal than without a meal. No main effect of statement type (both p’s > .237) nor interaction of nutritional state and statement type was observed for right or left VS (both p’s > .079). In the vmPFC ROI, brain activity was significantly higher for compliments than for neutral statements (F(1,51)=46.29, p<.001, η_p_^2^=.48), while nutritional state did not significantly influence vmPFC activation (both p’s > .571; see Table 4 and Figure 6).

**Figure 6.**
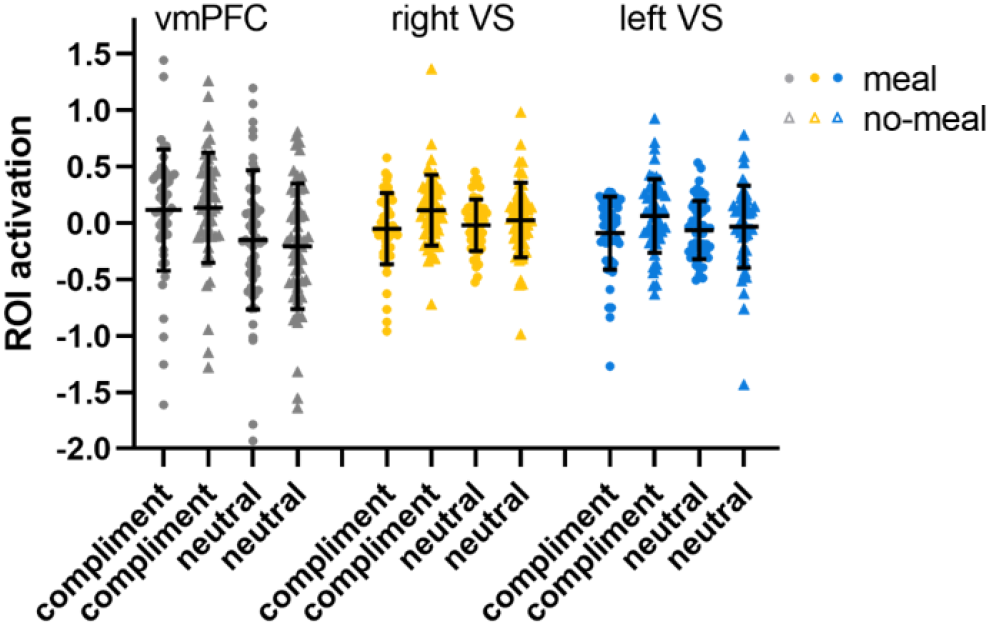
ROI activation in the social affirmation task. Black horizontal bars represent mean values, error bars represent 1 standard deviation. Single-subject data are plotted as dots (liquid-meal condition) or triangles (no-meal condition) for compliments and neutral statements. Please note that outlier values are included in this figure.

### 3.7 Task 2: Relationship between ghrelin concentrations at T1, pleasantness ratings and brain activation during compliments and neutral statements

Correlation analyses failed to show any significant associations between the ghrelin satiation response (Δghrelin at T1) and Δpleasantness ratings (both p’s > .564), and between Δghrelin at T1 and ΔROI brain activation (all p’s > .234). Regarding brain-behaviour correlations, a positive association between Δpleasantness ratings and ΔrightVS for compliments was observed (r_s_(50)=.336, p=.017). The higher the brain activation in right VS was during liquid-meal compared to no-meal, the higher were the pleasantness ratings in the liquid-meal compared to the no-meal session. No other analysis showed significant associations (all p-values > .048).

Bayesian analyses showed moderate evidence for no association between Δghrelin and Δpleasantness ratings following compliments and neutral statements, and between Δghrelin and ΔROI activation during compliments and neutral statements (all BF01>3). For the association between Δghrelin and ΔvmPFC activation during neutral statements, only anecdotal evidence was available for a null effect; see Table 6.

**Table 6.**
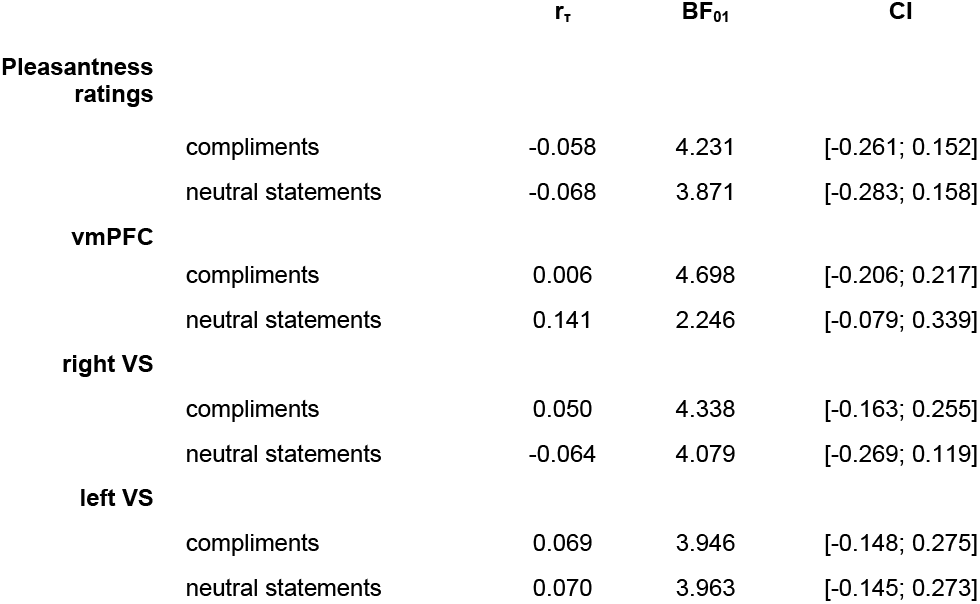

## 4. Discussion

This study investigated whether reward responses to social and nonsocial rewards are influenced by variation in the gut hormone ghrelin. The same fasting participants either received a meal to decrease their endogenous ghrelin concentrations or they had to continue fasting to maintain high ghrelin concentrations throughout the experiment. Contrary to the preregistered hypotheses, no significant association was observed between natural ghrelin variation and behavioural or neural markers of social reward processing. Activation in the right ventral striatum was higher when fasting than when having eaten during the presentation of both compliments and neutral statements, but nutritional state did not affect pleasantness. Additional Bayes analyses provided moderate evidence that ghrelin concentrations were not linked to behavioral or neural outcomes of social reward processing in both tasks. In an exploratory analysis of non-social rewards in task 1, larger ghrelin suppression after the meal was associated with larger suppression of vmPFC activity after the meal.

### 4.1 No association between ghrelin concentrations and the processing of social rewards

Our study did not show an association between ghrelin variation and social rewards. This is in contrast to previous studies with rodents (24,25) and healthy humans (27). However, at closer view, there are also alternative interpretations for the findings in these studies. For instance, mice with blocked or abolished ghrelin receptor signalling took longer time to approach a stranger mouse, and ghrelin receptor knock-out prevented this from happening (25). This could suggest that ghrelin receptor signalling promotes social behaviour, but it could also suggest that activation of this receptor promotes general motivation/exploration and that the only salient stimuli at this time were the novel conspecifics in the middle of the box. Thus, the observed behavioural changes could reflect salience or general motivation without a specific social component. In addition, ghrelin has also been linked to aggressive behaviours in mice (e.g., (51,52)), therefore even if ghrelin increases social approach, this may not lead to pro-social or positive social interactions.

A different explanation for the lack of association between ghrelin concentrations and the processing of social rewards in the present study could lie in the nature of the social rewards. The social rewards used in the previous studies and the present study differ in several aspects, namely being primary or secondary rewards, and being tangible or not (i.e., being touchable and consumable vs. being abstract), see (53) for an overview. Primary rewards are assumed to be innately rewarding, e.g. food or touch. Indeed, the rodent studies used a primary and tangible social reward, namely natural interaction with another conspecific that was present in the same space (24,25). Similarly, a human study providing first evidence for a potential role of the ghrelin system in social reward (27) used affective touch, i.e. caress-like gentle touch that is assumed to be innately rewarding (54,55).

In contrast, social rewards in task 1 and 2 constituted of secondary rewards, namely social appreciation and affirmation in written form on the screen. These rewards have to be learned from and interpreted in their respective social context. For example, depending on one’s cultural upbringing, social appreciation by unknown experts might be less rewarding if one values collective appreciation more than an individualistic one (56,57). In addition to being secondary, i.e. learned, this form of reward is also less tangible and more abstract then an approval in the form of a tap on the shoulder. Future studies are needed to clarify ghrelin’s role in the processing of these different instances of social rewards. Such abstract forms of social recognition and affirmation might be too complex to be susceptible to the influence of ghrelin.

### 4.2 Association between ghrelin suppression and vmPFC activation to non-social reward (exploratory analysis)

In contrast to the two forms of social reward, ghrelin variation was associated with non-social reward. A larger ghrelin suppression after the meal (i.e., a stronger ghrelin satiety response) was associated with a larger suppression of vmPFC activity after the meal for the non-social computer reward in task 1. If participants experienced only a small or even an absent ghrelin suppression after the meal compared to no meal, vmPFC activity was enhanced after the meal.

The vmPFC is a neural hub involved, among others, in the attribution of subjective relevance to surrounding stimuli and behaviours (58,59). In light of this, it appears as if the non-social computer rewards were valued differently depending on whether participants showed a large ghrelin suppression by the meal or not.

If ghrelin concentrations stayed on a heightened level individually, they seemed to be associated with enhanced vmPFC activation. This observation is in line with recent studies were high ghrelin concentrations predicted a preference for immediate (but smaller) monetary rewards in healthy individuals (21) as well as a preference for gambling persistence when confronted with losses (22). High ghrelin concentrations thus seemed to support perseverance behaviour during the anticipation of immediate monetary rewards in these studies. The same logic might apply to the current study. Participants with higher ghrelin concentrations after the meal might have anticipated the receipt of the USB stick with their favourite songs to a stronger degree than participants with a larger ghrelin suppression by the meal. This would further suggest that the ghrelin system might play a larger role during the anticipation than the consumption of rewards (60). This suggestion would be in line with studies reporting increased ghrelin secretion in anticipation of scheduled food intake (7,61).

The size of the meal-induced ghrelin suppression varied considerably across the sample. One possible explanation could be that participants with a smaller suppression might have needed more food for a stronger ghrelin decrease. Alternatively, participants’ genetic makeup could have influenced the size of the meal-induced ghrelin suppression. A recent study investigated a polymorphism in the fat mass and obesity-associated gene *FTO* and found that homozygous AA allele carries showed a smaller meal-induced ghrelin suppression after the same caloric load than TT allele carriers (62). Future studies could manipulate endogenous ghrelin concentrations in a more individualized way based on participants’ sex and body mass index and take the *FTO* polymorphism into account. Thereby, it could be systematically investigated whether the observed diverging effects of small vs. large ghrelin suppression on reward valuation replicate in, for example, food and non-food rewards.

### 4.3 Impact of nutritional state

Manipulating nutritional state by either providing a meal or keeping participants fasted showed the intended effects on ghrelin variation and subjective bodily experiences. At similar time points in the experiment, ghrelin concentrations were considerably reduced after a meal compared to no-meal, and subjective hunger ratings and participants’ desire to eat were higher without the meal. Participants reported slightly enhanced negative affect in the no-meal compared to the liquid-meal session, however with a small effect size and on the lower end of the assessment scale. This suggests that the current participants were rather not “hangry” (63) during the experiment, which could have otherwise influenced the processing of social rewards. No significant effects of nutritional state were observed in the social-recognition-by-experts task (task 1) for both behavioural and neural measures. While subjective pleasantness was also not influenced by nutritional state in the social affirmation task (task 2), brain activation in response to all types of statements was. In particular, right ventral striatum activation was higher when participants had not eaten compared to when they had, irrespective of whether compliments or neutral statements were presented. Descriptively, a similar activation pattern was observed in left ventral striatum but this did not reach significance.

Ventral striatum activation has repeatedly been associated with reward processing (59,64–66), but also more generally with stimulus salience attribution in the context of reward and punishment (67). Thus, cautiously interpreting the observed right ventral striatum activation, it seems that all types of statements were processed more strongly - and thus more rewarding or salient - when participants had not eaten compared to after a meal. Both compliments and neutral statements referred to the participants themselves, which may have increased self-directed attention, but may have also signalled increased attention from others towards the participants. A previous study found selective activation to the passive viewing of food cues after fasting and to social cues after isolation in midbrain regions, but not in the striatum (68). Thus, an effect of fasting on striatal activation may differ between passive picture viewing and the presentation of statements which may signal social attention. Likewise, we have previously observed that perceived social isolation is associated with altered striatal responses to social feedback videos which also reflect social attention (69).

### 4.4 Limitations

The current within-subject feed-and-fast approach allowed only an indirect manipulation of endogenous ghrelin concentrations. Thus, no causal relationships (or the absence thereof) between ghrelin concentrations and the assessed outcome measures can be inferred. However, the current approach provided the advantage of investigating intra-individual ghrelin variation within a physiologically plausible range, which allows for higher ecological validity. Our approach is in stark contrast to studies utilizing intravenous ghrelin administration leading to short-lasting artificially high ghrelin concentrations.

Ghrelin concentrations were only assessed three times during the experiment. An additional assessment shortly after consuming the snack before the scanning session (and at the corresponding time point in the no-meal session) would have provided additional information on continued ghrelin suppression, but no such sample was taken. The current T1 measurement was taken at least 30 min before starting the scanner, so it should be considered an approximation of true ghrelin concentrations during scanning. Future studies should aim at implementing continuous ghrelin sampling throughout the whole experimental procedure.

## 5. Conclusion

Naturally circulating ghrelin concentrations were not associated with behavioural or neural responses to social rewards, but to responses to non-social rewards. This could be due to differences in the social nature of rewards or their tangibility. Alternatively, ghrelin may affect the anticipation of reward rather than the response to its consumption.

## Data availability

All data, code, and materials will be made publicly available via the Open Science Framework: ***

The study was pre-registered on the Open Science Forum (https://osf.io/f9rkq). This manuscript was deposited as a preprint on the server bioRxiv *** under a CC-BY-NC-ND 4.0 International license.

## Acknowledgements

We thank TINE BA for providing us with the products used as liquid meal for free. We thank Erik R. Frogner, Aiste Gvildyte Næss, Pietro Aleksander Rocco Berger Lapolla, Aleksandra Pusica, Thea Wiker Engelund, and Anbjørn Ree for their work with participant recruitment and data collection.

## Author contributions

Original idea: Uta Sailer; Designed the study: Uta Sailer, Daniela M. Pfabigan; Provided input to methods: Daniel Campbell-Meiklejohn; Performed the research: Daniela M. Pfabigan; Data curation: Daniela M. Pfabigan, Federica Riva; Data analyses: Federica Riva, Daniela M. Pfabigan; Data visualization: Federica Riva, Daniela M. Pfabigan; Wrote the manuscript: Uta Sailer, Daniela M. Pfabigan, Federica Riva; Provided comments on the manuscript: Jana Lieberz, Dirk Scheele, Daniel Campbell-Meiklejohn; Acquired funding: Uta Sailer. All authors contributed and agreed to the final version of the manuscript.

## Footnotes

1 The reverse contrasts (disagreement > agreement; no-win > win; FWE peak- or cluster-level correction, tested in the liquid-meal session) did not result in significant activation clusters.

